# Human Attitudes Toward Insects and Spiders: Exploring the Paradoxes of Ecological Value and Discomfort

**DOI:** 10.1101/2025.07.31.667848

**Authors:** Divya Arora, Sunil Dutt Shukla

## Abstract

Insects, with over a million species, are important to ecosystems, contributing to pollination, nutrient cycling, and pest control, yet public perceptions often lean negative due to fear, disgust, or cultural influences. The study explores human attitudes toward insects via a 17-question Google Forms survey, finding consistent perceptions across demographics, except for education level, which impacts ecological awareness and tolerance. A notable exception was the belief that all insects are harmful, which varied significantly by age, and gender differences in handling dangerous insects. Principal Component Analysis identified three key dimensions—environmental awareness, demographic influences, and emotional responses— explaining 38% of the variance. The findings highlight paradoxes in public views, such as recognizing insects’ ecological value while expressing discomfort with their presence. Education emerges as a critical factor in fostering positive attitudes, suggesting targeted campaigns could bridge knowledge gaps and promote coexistence with these essential creatures.

## Introduction

Insects comprise over a million species, playing crucial roles in ecosystems like pollination and nutrient recycling. This diversity maximum in among any taxonomic groups, demonstrating the success of insects in evolution. They inhabit in nearly all ecosystems, ranging from tropical rainforests to arid deserts, and exhibit a surprising range of adaptions (morphological, behavioural, and ecological). This diversity underscores their important role in global biodiversity and emphasizes the need to preserve insect communities (Free, 1970; Pellmyr, 1992). Insects play role in nutrient cycling by decomposing organic matter (Abajue & Gbarakoro, 2023; Verma et al., 2023). Dung beetles, for example, contribute to soil enrichment by burying and decomposing animal feces. In addition to nutrient cycling, many insects act as natural pest controllers. Apart from nutrient cycling, most insects play the role of natural pest control. Predatory insects like ladybugs and lacewings prey on agricultural pests, thus lowering the usage of chemical pesticides. Current declines in insect populations, such as important pollinators such as bees and charismatic pollinators like butterflies, are major threats to biodiversity and ecosystem stability (Abajue & Gbarakoro, 2023; Barragán-Fonseca, 2024; Cadinu et al., 2020). Insect-human interaction goes thousands of years back, with arthropods having adapted to human habitation and domestic settings. Archaeological evidence and records suggests the close association between humans and insects; began when early humans inhibited caves, forming stable, sheltered microhabitats for some arthropods (Berenbaum, 1996; Jose, 2019; Ramos-Elorduy, 2009). Despite their importance, many individuals do feel negative towards insects, including fear and disgust, often due to cultural factors, ignorance, or past experiences (Fukano & Soga, 2023; Soga et al., 2023).

Perceptions of insects are diverse across cultures, from symbols of transformation and renewal to pests. In various Asian and African cultures, entomophagy—the practice of eating insects— is entranced in traditional diets and often act as a delicacy for its nutritional, economic, and environmental reasons. For example, In parts of Thailand, Zimbabwe, and Nigeria, for instance, insects such as crickets, locusts, and caterpillars are commonly eaten and enjoyed as an sustainable source of protein (Ramos-Elorduy, 2009).

Positive experiences and education can change attitudes, highlighting insects’ vital contributions in ecosystems, such as pollination and decomposition (Gaston, 1991; Pellmyr, 1992). Negative feelings towards insects are defined as entomophobia or disgust, frequently associated to their appearance or being seen as vector for disease (Fukano & Soga, 2023).

The media is a strong influencer of the public perception of insects, often perpetuating negative typecasts by portraying them as pests, threats, or sources of fear. Depictions in movies, television shows, and advertisements highlights the annoyance or danger associated with insects, e.g., infestations, bites, and the disease transmission (Cassel, 2016; Snaddon & Turner, 2007).

However, Human also developed a more positive outlook on insects by recognizing their importance, understanding their role in nature, and supporting conservation (Infield et al., 2018). Soga et al., (2023) suggest that having people around you who work for nature conservation may enhance positive attitudes towards insects, this encourages curiosity and appreciation towards the important roles insects play. This all is helpful in better understanding and conservation of these essential creatures.

As Berenbaum (1996) records, "Like it or not, insects are a part of where we have come from, what we are now, and what we will be." We can promote biodiversity conservation efforts and value their role in shaping our environment and culture by acknowledging their environmental significance (Yadav et al., 2023).

Present study was grounded in the recognition that insects play a vital ecological role, yet often elicit strong emotional responses ranging from fascination to fear or disgust. Understanding these perceptions is essential to promote awareness, conservation, and coexistence with insects, especially in the face of biodiversity loss.

## Material and Methods

The objective of this study was to assess human attitudes and perception toward insects and how these attitudes vary across demographic categories like gender, educational qualification, and age group.

To achieve this, a cross-sectional survey design was employed, which made it possible to collect quantitative data at a single point in time. A structured questionnaire was developed to assess participants’ knowledge, attitudes, and perceptions of insects in both every day and ecological contexts. The questionnaire comprised 17 questions (Table 1), designed in multiple-choice and Likert-scale formats.

**Table 1:** Survey Questions on People’s Perception of Insects by Category.

| Category | Questions |
| --- | --- |
| Demographic Information | 1. Your Name<br>2. Age<br>3. Gender<br>4. Location<br>5. Education/Qualification |
| General Perception of Insects | 6. Do you generally like insects?<br>7. Do you consider insects to be helpful to the environment?<br>8. When you see an insect, is your first reaction to feel scared and disgusted?<br>9. Do you believe that all insects are harmful to humans? |
| Emotional Reaction and Attitudes | 10. How often do you feel uncomfortable when you encounter an insect in your living space?<br>11. Which of the following types of insects do you find most attractive?<br>12. Which of the following types of insects do you find most disturbing?<br>13. If an insect lands on your skin, how would you react? |
| Awareness and Knowledge | 14. Are you aware that insects play a crucial role in pollination, decomposition, and food chain?<br>15. Do you think insects should be protected due to their ecological importance? |
| Insects Control and Interaction | 16. Have you ever used chemical products (e.g., insecticide) to kill insects?<br>17. If you encounter a dangerous insect (e.g., venomous spider, aggressive wasp), do you believe it's best to kill it immediately?<br>18. Do you think that insects should be removed from your home, or should they be left alone if they are not dangerous? |
| Personal Beliefs and Preferences | 19. Which of the following best describes your opinion on insects?<br>20. Would you feel comfortable living in a place where insects are allowed to freely roam indoors? |
| Future Consideration | 21. Would you be open to eating insects as a source of protein in the future?<br>22. Do you think insects will play a major role in addressing environmental or food security issues in the future? |

The questions were on various aspects such General perception, Emotional Response, Attractiveness and Disturbance, Environmental Importance and Protection, dealing with insects and future perspective. The survey was administered online using Google Forms, facilitating efficient data collection (Supplementary Data 1).

Participants were recruited using a convenience sampling method, targeting a diverse population across urban and semi-urban areas, genders, ages and was disseminated via email and various social media platforms over two weeks.

Given the nature of this study, which involved a voluntary, anonymous online survey with informed consent, ethical committee approval was not sought as no personal or sensitive information was collected during the study.

The responses were recorded onto a Google Sheet, where data anonymity was maintained. Only aggregate data entered Excel and JASP for further analysis.

To evaluate differences in insect perception across demographic categories, non-parametric statistical tests were employed, considering the ordinal nature of the data and the relatively small sample sizes in certain subgroups. Specifically, the Chi-Square Test for Independence (χ² test) was used to examine associations between categorical variables, such as age group, gender, and residential status, with responses to individual questionnaire items. This test helped determine whether the observed differences in response frequencies across demographic groups were statistically significant. A significance level of α = 0.05 was applied throughout the analysis, and the corresponding degrees of freedom (df) and chi-square values (χ²) were reported for each test to assess the strength and reliability of associations.

In addition to inferential analysis, descriptive statistics—namely, means and standard deviations (Mean±SD)—were calculated for each group to summarize central tendencies and the spread of responses.

Study purpose was well conveyed to participants and participation was entirely voluntary in nature. Participants were informed of the purpose of the study and that it was completely voluntary in nature. Anonymity was maintained by avoiding collecting any identifiable details (contact number, mail id etc).

## Results

The study provides an understanding of how people perceive insects; it also concentrates on the interplay between demographic factors. The study recognises areas of commonality and divergence in attitudes toward insects. The study highlights the importance of education in shaping perceptions, stating that higher education levels associated with greater awareness of ecological importance of insects. Eventually guiding initiatives towards appreciation and coexistence with these essential creatures.

Across all demographics, there is near-universal agreement (90–100%) on insects’ ecological importance, as evidenced by responses to Q2, Q9, Q10, and Q17. Discomfort with insects in living spaces (Q5, Q15) and fear reactions (Q3) are common, particularly among lower-educated (XII), urban, and middle-aged (21–30, 31–40) respondents. Demographic differences highlight nuanced patterns: females and the 21–30 group are more likely to kill dangerous insects, while males, rural residents, and younger/lower-educated respondents show slightly more openness to eating insects (8–23%, Q16). Higher education correlates with positive attitudes, less fear, and greater tolerance for non-dangerous insects. Butterflies are universally seen as attractive, while mosquitoes and cockroaches are most disturbing. These findings suggest that while ecological awareness is high, emotional and behavioural responses to insects vary by demographic, with implications for education and policy to promote insect conservation and acceptance. (Supplementary Data 2-4, contingency tables).

### Insect Perception Responses by Age Group (Table-2)

The study on insect perception across different age groups revealed no significant differences in responses for any of the 17 questions. This suggests that age does not play a substantial role in shaping perceptions towards insects.

**Table 2:** Insect Perception Responses by Age Group (Valus depicted as Mean±SD and *Significant at p < 0.05 (chi-square test))

| Q.No | Below 20 | 21–30 | 31–40 | 41–50 | Above 50 | $\chi^2$ | df | p-value |
| --- | --- | --- | --- | --- | --- | --- | --- | --- |
| 1 | 1.54±0.52 | 1.50±0.51 | 1.33±0.58 | 1.67±0.58 | 1.75±0.46 | 2.323 | 4 | 0.677 |
| 2 | 1.83±0.39 | 1.97±0.18 | 2.00±0.00 | 2.00±0.00 | 2.00±0.00 | 4.062 | 4 | 0.398 |
| 3 | 1.46±0.52 | 1.53±0.51 | 1.67±0.58 | 1.33±0.58 | 1.38±0.52 | 1.360 | 4 | 0.851 |
| 4 | 1.08±0.28 | 1.07±0.25 | 1.33±0.58 | 1.00±0.00 | 1.00±0.00 | 4.029 | 4 | 0.402 |
| 5 | 1.62±0.87 | 1.83±0.83 | 2.67±1.53 | 2.00±1.73 | 2.00±1.31 | 14.708 | 12 | 0.258 |
| 6 | 1.92±1.24 | 1.83±0.95 | 2.00±1.00 | 2.67±0.58 | 2.88±1.55 | 83.316 | 68 | 0.100 |
| 7 | 1.64±0.92 | 2.03±1.03 | 2.67±1.53 | 2.67±1.53 | 2.25±1.04 | 90.055 | 96 | 0.652 |
| 8 | 2.69±0.63 | 2.33±0.66 | 2.33±0.58 | 2.67±0.58 | 2.75±0.46 | 7.541 | 8 | 0.480 |
| 9 | 1.92±0.29 | 1.97±0.18 | 2.00±0.00 | 2.00±0.00 | 2.00±0.00 | 1.314 | 4 | 0.859 |
| 10 | 2.00±0.29 | 2.00±0.00 | 2.00±0.00 | 2.00±0.00 | 2.00±0.00 | 3.733 | 4 | 0.443 |
| 11 | 1.36±0.51 | 1.47±0.51 | 1.67±0.58 | 1.33±0.58 | 1.62±0.52 | 1.960 | 4 | 0.743 |
| 12 | 1.46±0.52 | 1.67±0.48 | 1.33±0.58 | 1.67±0.58 | 1.25±0.46 | 5.682 | 4 | 0.224 |
| 13 | 1.33±0.49 | 1.47±0.51 | 1.33±0.58 | 1.33±0.58 | 1.50±0.54 | 1.011 | 4 | 0.908 |
| 14 | 2.31±1.44 | 2.00±1.05 | 2.00±1.73 | 1.67±0.58 | 1.25±0.46 | 12.666 | 12 | 0.394 |
| 15 | 1.46±0.52 | 1.13±0.35 | 1.00±0.00 | 1.00±0.00 | 1.12±0.35 | 8.378 | 4 | 0.079 |
| 16 | 1.23±0.44 | 1.17±0.38 | 1.00±0.00 | 1.00±0.00 | 1.00±0.00 | 3.339 | 4 | 0.503 |
| 17 | 1.92±0.28 | 1.90±0.31 | 2.00±0.00 | 2.00±0.00 | 2.00±0.00 | 1.475 | 4 | 0.831 |

Respondents generally held positive views about the environmental role of insects. For instance, when asked if insects are helpful to the environment, mean scores ranged from 1.83 to 2.00 across age groups, indicating a strong agreement (χ² = 4.062, p = 0.398). Similarly, awareness of the ecological role of insects was high, with mean scores ranging from 1.92 to 2.00 (χ² = 1.314, p = 0.859).

However, despite recognizing the importance of insects, respondents expressed discomfort and fear towards them. When asked if they were scared or disgusted by insects, mean scores ranged from 1.33 to 1.67 (χ² = 1.360, p = 0.851), and when asked about comfort levels with insects in their living space (Q5), mean scores ranged from 1.62 to 2.67 (χ² = 14.708, p = 0.258). This ambivalence was also reflected in their willingness to protect insects, with mean scores of 2.00 across all age groups (χ² = 3.733, p = 0.443), while also being less likely to use insecticides or kill insects unless necessary.

Interestingly, respondents were generally not open to eating insects, with mean scores ranging from 1.00 to 1.23 (χ² = 3.339, p = 0.503). Overall, the findings highlight the complexity of human-insect relationships, with people recognizing the importance of insects while also harbouring negative emotions towards them. These insights can inform strategies for promoting insect conservation and mitigating conflicts between humans and insects.

### Insect Perception Responses by Gender (Table-3)

The analysis of insect perception responses by gender shows that overall attitudes between females and males are largely similar, with only one statistically significant difference observed. In general, both females (mean = 1.50) and males (mean = 1.59) showed relatively low levels of liking insects, with no significant difference (χ² = 0.491, p = 0.483). Both genders also strongly agreed that insects are helpful to the environment (mean = 1.97 for females, 1.92 for males), and displayed similar levels of ecological awareness, including agreement on insects’ roles in pollination and the food chain.

**Table 3:** Insect Perception Responses by Gender (Valus depicted as Mean±SD and *Significant at p < 0.05 (chi-square test))

| **Q. No** | **Female** | **Male** | **$\chi^2$** | **df** | **p** |
| --- | --- | --- | --- | --- | --- |
| 1 | 1.50±0.51 | 1.59±0.50 | 0.491 | 1 | 0.483 |
| 2 | 1.97±0.18 | 1.92±0.27 | 0.522 | 1 | 0.470 |
| 3 | 1.47±0.51 | 1.52±0.51 | 0.153 | 1 | 0.696 |
| 4 | 1.07±0.25 | 1.07±0.27 | 0.012 | 1 | 0.913 |
| 5 | 1.97±0.93 | 1.74±1.06 | 7.619 | 3 | 0.055 |
| 6 | 2.23±1.10 | 1.85±1.16 | 17.188 | 17 | 0.442 |
| 7 | 2.27±1.02 | 1.80±1.08 | 24.891 | 24 | 0.412 |
| 8 | 2.53±0.57 | 2.44±0.70 | 1.400 | 2 | 0.497 |
| 9 | 1.97±0.18 | 1.96±0.20 | 0.011 | 1 | 0.918 |
| 10 | 2.00±0.00 | 1.96±0.20 | 1.175 | 1 | 0.278 |
| 11 | 1.50±0.51 | 1.44±0.51 | 0.197 | 1 | 0.657 |
| 12 | 1.70±0.47 | 1.37±0.49 | 6.224 | 1 | 0.013* |
| 13 | 1.37±0.49 | 1.50±0.51 | 1.011 | 1 | 0.315 |
| 14 | 1.83±1.02 | 2.07±1.24 | 2.294 | 3 | 0.514 |
| 15 | 1.17±0.38 | 1.22±0.42 | 0.282 | 1 | 0.596 |
| 16 | 1.07±0.25 | 1.22±0.42 | 2.850 | 1 | 0.091 |
| 17 | 1.90±0.31 | 1.96±0.19 | 0.863 | 1 | 0.353 |

There was little difference in emotional reactions to insects, such as feeling scared or disgusted, or in the belief that all insects are harmful—both genders strongly disagreed with that notion (mean = 1.07 for both). Female respondents reported slightly more discomfort when encountering insects indoors (mean = 1.97) compared to males (mean = 1.74), approaching but not reaching statistical significance (χ² = 7.619, p = 0.055).

Preferences in finding insects attractive or disturbing showed variation but were not statistically significant (p = 0.442 and p = 0.412, respectively). Reactions to insects landing on the skin, use of chemical products, and comfort with insects indoors also showed no meaningful gender-based differences.

However, a significant gender difference emerged in the belief that dangerous insects (e.g., venomous spiders or aggressive wasps) should be killed immediately. Females (mean = 1.70) were more likely to support killing them compared to males (mean = 1.37), and this difference was statistically significant (χ² = 6.224, p = 0.013), indicating a more defensive stance among women.

On the topic of entomophagy (eating insects), females were less open (mean = 1.07) than males (mean = 1.22), though this difference did not reach statistical significance (χ² = 2.850, p = 0.091). Similarly, opinions on the future role of insects in addressing environmental and food security issues showed no significant gender gap.

### Insect Perception Responses by Resident Status (Table-4)

The survey data comparing insect perception between rural and urban residents revealed generally similar attitudes with only minor differences, none of which were statistically significant. When asked whether they generally like insects, both rural (mean = 1.57) and urban (mean = 1.53) participants responded similarly, indicating a general dislike or ambivalence, with no significant difference (χ² = 0.057, p = 0.812). Likewise, both groups strongly agreed that insects are helpful to the environment, although rural residents unanimously agreed (mean = 2.00), while a small variation was observed among urban respondents (mean = 1.93), still without a significant statistical difference (χ² = 1.057, p = 0.304).

**Table 4:** Insect Perception Responses by Resident Status (Valus depicted as Mean±SD and *Significant at p < 0.05 (chi-square test))

| Q. No | Rural | Urban | $\chi^2$ | df | p |
| --- | --- | --- | --- | --- | --- |
| 1 | 1.57±0.51 | 1.54±0.51 | 0.057 | 1 | 0.812 |
| 2 | 2.00±0.00 | 1.93±0.26 | 1.057 | 1 | 0.304 |
| 3 | 1.50±0.52 | 1.49±0.51 | 0.006 | 1 | 0.940 |
| 4 | 1.07±0.27 | 1.07±0.26 | 0.00 | 1 | 0.983 |
| 5 | 2.36±1.15 | 1.70±0.89 | 5.499 | 3 | 0.139 |
| 6 | 1.79±1.05 | 2.14±1.16 | 15.883 | 17 | 0.532 |
| 7 | 2.14±1.17 | 2.02±1.04 | 23.276 | 24 | 0.504 |
| 8 | 2.50±0.65 | 2.49±0.63 | 0.010 | 2 | 0.995 |
| 9 | 2.00±0.00 | 1.95±0.22 | 0.691 | 1 | 0.406 |
| 10 | 2.00±0.00 | 1.98±0.15 | 0.339 | 1 | 0.560 |
| 11 | 1.43±0.51 | 1.49±0.51 | 0.147 | 1 | 0.702 |
| 12 | 1.50±0.52 | 1.56±0.50 | 0.144 | 1 | 0.704 |
| 13 | 1.43±0.51 | 1.43±0.50 | 0.00 | 1 | 1.00 |
| 14 | 1.86±1.03 | 1.98±1.17 | 1.103 | 3 | 0.776 |
| 15 | 1.14±0.36 | 1.21±0.41 | 0.299 | 1 | 0.584 |
| 16 | 1.21±0.43 | 1.12±0.32 | 0.841 | 1 | 0.359 |
| 17 | 2.00±0.00 | 1.91±0.29 | 1.401 | 1 | 0.237 |

Initial emotional reactions such as fear or disgust upon seeing an insect were comparable across the two groups (mean = 1.50 rural vs. 1.49 urban; χ² = 0.006, p = 0.940). Similarly, both rural and urban participants strongly disagreed with the statement that all insects are harmful to humans, with identical mean scores (1.07) and no significant difference. The question about discomfort with insects indoors showed a higher mean for rural respondents (2.36) than urban respondents (1.70), suggesting greater discomfort among rural individuals, though this difference was still not statistically significant (χ² = 5.499, p = 0.139).

Preferences for attractive or disturbing insects varied slightly between groups but did not show any significant patterns, with p-values of 0.532 and 0.504 respectively. Reactions to an insect landing on the skin were nearly identical (mean = 2.50 rural vs. 2.49 urban), and awareness of insects’ ecological roles was almost universal, especially among rural respondents, who again scored a full 2.00 mean.

When asked whether insects should be protected due to ecological importance, rural participants unanimously agreed (mean = 2.00), with urban residents showing near-unanimous agreement (mean = 1.98), and no significant difference observed (χ² = 0.339, p = 0.560). The use of insecticides, beliefs about killing dangerous insects, and attitudes towards removing insects from the home all showed minimal variation between rural and urban residents, with no statistical significance.

Questions regarding comfort with insects roaming indoors, openness to eating insects as a protein source, and beliefs about the future role of insects in solving environmental or food security problems all yielded similar responses between rural and urban groups, with none showing significant differences.

### Insect Perception Responses by Qualification (Table-5)

The survey examined insect perception across different educational qualifications, ranging from Class X and XII students to individuals with postgraduate and doctoral degrees. Overall, participants generally did not express a strong liking for insects, though the degree of aversion varied slightly by educational level. Class XII respondents showed the least liking (mean = 1.20), while the "Other" category, with a single response, showed complete neutrality or indifference (mean = 2.00). However, responses to whether insects are helpful to the environment revealed a statistically significant difference across groups (χ² = 14.279, p = 0.014), indicating that individuals with higher education levels tend to be more aware of the ecological importance of insects. This trend is further supported by high awareness among postgraduate and PhD holders that insects contribute to pollination, decomposition, and food chains.

**Table 5:** Insect Perception Responses by Qualification (Valus depicted as Mean±SD and *Significant at p < 0.05 (chi-square test))

| Q. No. | X | XII | Graduation | Post Graduation | Ph D | Other | $\chi^2$ | df | p |
| --- | --- | --- | --- | --- | --- | --- | --- | --- | --- |
| 1. | 1.714 ± 0.488 | 1.200 ± 0.447 | 1.600 ± 0.516 | 1.571 ± 0.507 | 1.417 ± 0.515 | 2.000 ± 0.000 | 4.987 | 5 | 0.417 |
| 2. | 2.000 ± 0.000 | 1.600 ± 0.548 | 1.900 ± 0.316 | 2.000 ± 0.000 | 2.000 ± 0.000 | 2.000 ± 0.000 | 14.279 | 5 | 0.014* |
| 3. | 1.143 ± 0.378 | 1.800 ± 0.447 | 1.600 ± 0.516 | 1.476 ± 0.512 | 1.500 ± 0.522 | 1.000 ± 0.000 | 6.756 | 5 | 0.239 |
| 4. | 1.000 ± 0.000 | 1.200 ± 0.447 | 1.200 ± 0.422 | 1.000 ± 0.000 | 1.083 ± 0.289 | 1.000 ± 0.000 | 5.995 | 5 | 0.307 |
| 5. | 1.714 ± 1.113 | 1.600 ± 0.548 | 1.600 ± 0.843 | 2.000 ± 0.949 | 2.167 ± 1.267 | 1.000 ± 0.894 | 11.479 | 15 | 0.718 |
| 6. | 2.000 ± 0.837 | 1.200 ± 1.095 | 1.900 ± 1.337 | 2.143 ± 0.949 | 2.250 ± 1.084 | 5.000 ± 0.488 | 112.794 | 85 | 0.024* |
| 7. | 1.500 ± 0.894 | 1.800 ± 0.707 | 2.300 ± 0.590 | 2.000 ± 0.492 | 2.417 ± 0.000 | — | — | — | — |
| 8. | 2.714 ± 0.447 | 2.400 ± 0.316 | 2.500 ± 0.000 | 2.381 ± 0.000 | 2.667 ± 0.000 | 3.000 ± 0.000 | 7.400 | 10 | 0.687 |
| 9. | 2.000 ± 0.000 | 1.800 ± 0.447 | 1.900 ± 0.000 | 2.000 ± 0.000 | 2.000 ± 0.000 | 2.000 ± 0.000 | 6.486 | 5 | 0.262 |
| 10. | 2.000 ± 0.000 | 1.800 ± 0.447 | 2.000 ± 0.000 | 2.000 ± 0.000 | 2.000 ± 0.000 | 2.000 ± 0.000 | 10.185 | 5 | 0.070 |
| 11. | 1.429 ± 0.488 | 1.500 ± 0.548 | 1.400 ± 0.483 | 1.429 ± 0.498 | 1.667 ± 0.492 | — | — | — | — |
| 12. | 1.714 ± 0.408 | 1.400 ± 0.548 | 1.700 ± 0.483 | 1.619 ± 0.483 | 1.333 ± 0.515 | — | 6.036 | 5 | 0.303 |
| 13. | 1.571 ± 1.397 | 1.750 ± 1.643 | 1.000 ± 1.135 | 1.571 ± 0.746 | 1.600 ± 1.138 | — | 6.844 | 5 | 0.233 |
| 14. | 1.143 ± 0.535 | 1.000 ± 0.548 | 1.000 ± 0.316 | 1.286 ± 0.359 | 1.200 ± 0.000 | — | 18.625 | 15 | 0.231 |
| 15. | 1.714 ± 0.488 | 1.200 ± 0.447 | 1.600 ± 0.422 | 1.619 ± 0.218 | 1.333 ± 0.289 | — | 15.544 | 5 | 0.008* |
| 16. | 1.000 ± 0.000 | 1.000 ± 0.447 | 1.000 ± 0.316 | 1.000 ± 0.218 | 1.000 ± 0.289 | — | 9.469 | 5 | 0.092 |
| 17. | 1.952 ± 0.535 | 1.917 ± 0.548 | 2.000 ± 0.316 | 2.000 ± 0.359 | 2.000 ± 0.000 | — | 2.190 | 5 | 0.822 |

Participants generally disagreed with the notion that all insects are harmful, with mean scores close to 1.00 across all groups, indicating a correct understanding. Reactions to insects, particularly feelings of fear or disgust when encountering one, were somewhat more pronounced in the lower qualification groups but did not show a statistically significant difference. When it came to preferences and aversions, the question about which insects were found most attractive showed significant variation among groups (χ² = 112.794, p = 0.024), suggesting differing attitudes and exposure to insect types across educational backgrounds.

Comfort with insects in indoor environments also varied significantly (χ² = 15.544, p = 0.008), with individuals in higher education levels being more tolerant, while those with lower qualifications reported higher discomfort. Similarly, responses to whether insects should be protected due to their ecological importance were generally supportive across all education levels, with marginal statistical significance (χ² = 10.185, p = 0.070), again indicating stronger ecological sensitivity among the more educated groups. Most respondents admitted to using chemical insecticides, but this did not vary much across groups, nor did the belief that dangerous insects should be killed immediately.

Interestingly, the willingness to eat insects as a protein source showed some reluctance across all levels, with slightly more openness among those with lower qualifications, though the difference was not statistically significant. Lastly, the belief in insects playing a role in addressing environmental or food security challenges in the future was nearly unanimous across all groups, with high mean scores and no significant differences.

### Principal Component Analysis (PCA) (Table-6)

A Principal Component Analysis (PCA) was conducted to explore the underlying structure of attitudes and beliefs about insects. The analysis revealed three components that together accounted for 37.9% of the total variance in the dataset. To this variance all three components contributed almost equally the share of first component (RC1), Environmental Awareness & Ecological Understanding, was 13.6%, this was followed by 12.7% of second component (RC2), Demographics and Preferences and third component (RC3), Emotional and Behavioural Reactions, accounted for 11.7% of the variance.

**Table 6:**
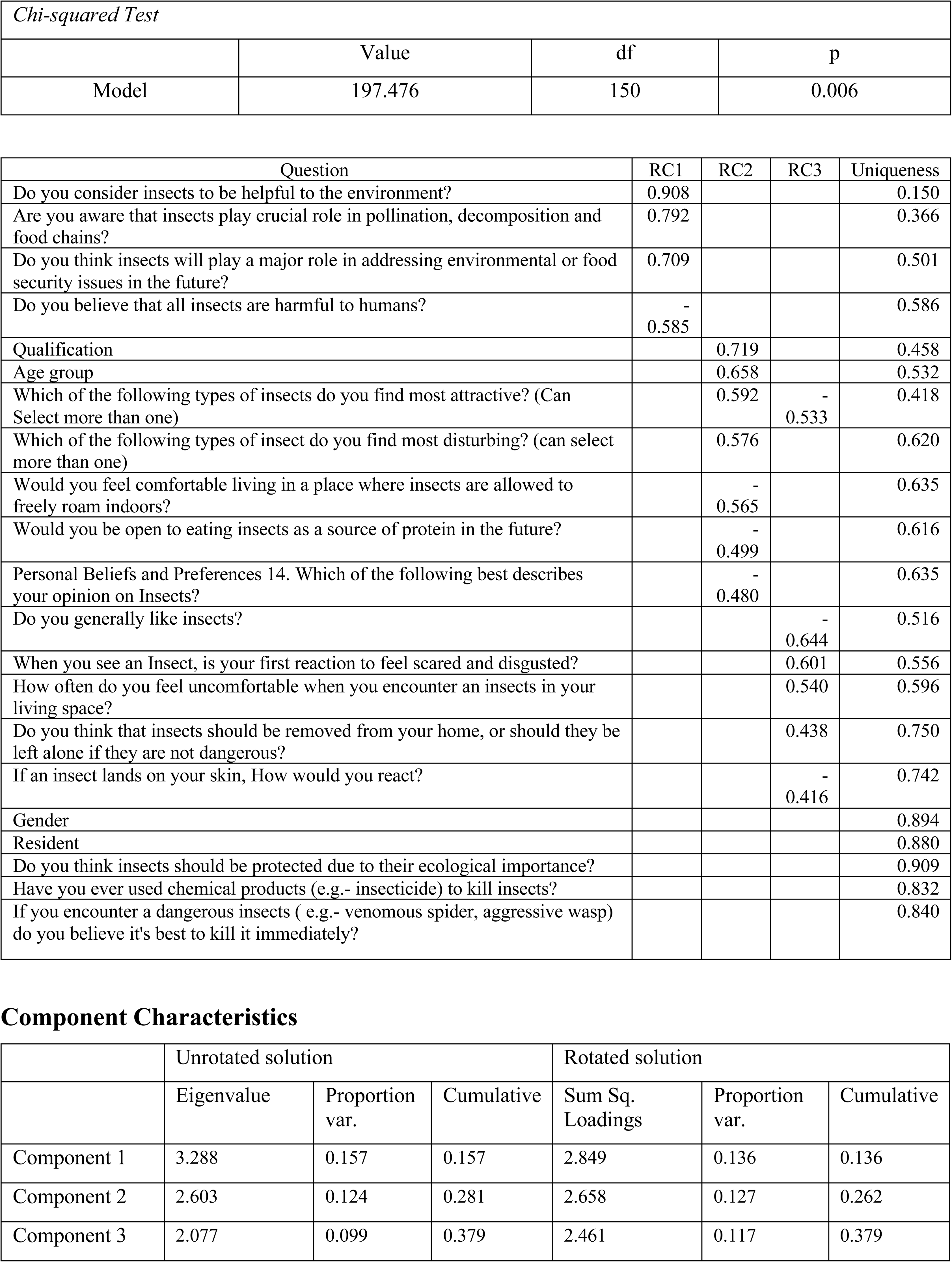
Principal Component Analysis of Insect Perception Responses. (RC1, RC2, and RC3 represent the three retained components, Values in the table represent factor loadings. Uniqueness values represent the proportion of variance in each variable not explained by the retained components. Eigenvalues and proportion of variance explained are presented for both unrotated and rotated solutions.)

| *Chi-squared Test* |  |  |  |
| --- | --- | --- | --- |
|  | Value | df | p |
| Model | 197.476 | 150 | 0.006 |

| Question | RC1 | RC2 | RC3 | Uniqueness |
| --- | --- | --- | --- | --- |
| Do you consider insects to be helpful to the environment? | 0.908 |  |  | 0.150 |
| Are you aware that insects play crucial role in pollination, decomposition and food chains? | 0.792 |  |  | 0.366 |
| Do you think insects will play a major role in addressing environmental or food security issues in the future? | 0.709 |  |  | 0.501 |
| Do you believe that all insects are harmful to humans? | -<br>0.585 |  |  | 0.586 |
| Qualification |  | 0.719 |  | 0.458 |
| Age group |  | 0.658 |  | 0.532 |
| Which of the following types of insects do you find most attractive? (Can Select more than one) |  | 0.592 | -<br>0.533 | 0.418 |
| Which of the following types of insect do you find most disturbing? (can select more than one) |  | 0.576 |  | 0.620 |
| Would you feel comfortable living in a place where insects are allowed to freely roam indoors? |  | -<br>0.565 |  | 0.635 |
| Would you be open to eating insects as a source of protein in the future? |  | -<br>0.499 |  | 0.616 |
| Personal Beliefs and Preferences 14. Which of the following best describes your opinion on Insects? |  | -<br>0.480 |  | 0.635 |
| Do you generally like insects? |  |  | -<br>0.644 | 0.516 |
| When you see an Insect, is your first reaction to feel scared and disgusted? |  |  | 0.601 | 0.556 |
| How often do you feel uncomfortable when you encounter an insects in your living space? |  |  | 0.540 | 0.596 |
| Do you think that insects should be removed from your home, or should they be left alone if they are not dangerous? |  |  | 0.438 | 0.750 |
| If an insect lands on your skin, How would you react? |  |  | -<br>0.416 | 0.742 |
| Gender |  |  |  | 0.894 |
| Resident |  |  |  | 0.880 |
| Do you think insects should be protected due to their ecological importance? |  |  |  | 0.909 |
| Have you ever used chemical products (e.g.- insecticide) to kill insects? |  |  |  | 0.832 |
| If you encounter a dangerous insects ( e.g.- venomous spider, aggressive wasp) do you believe it's best to kill it immediately? |  |  |  | 0.840 |

**Table 6: Principal Component Analysis of Insect Perception Responses** (RC1, RC2, and RC3 represent the three retained components, Values in the table represent factor loadings.
|  | Unrotated solution |  |  | Rotated solution |  |  |
| --- | --- | --- | --- | --- | --- | --- |
|  | Eigenvalue | Proportion var. | Cumulative | Sum Sq. Loadings | Proportion var. | Cumulative |
| Component 1 | 3.288 | 0.157 | 0.157 | 2.849 | 0.136 | 0.136 |
| Component 2 | 2.603 | 0.124 | 0.281 | 2.658 | 0.127 | 0.262 |
| Component 3 | 2.077 | 0.099 | 0.379 | 2.461 | 0.117 | 0.379 |

The first component (RC1) is strongly associated with environmental awareness and ecological understanding and exhibited positive attitudes toward insect. The second component (RC2), Demographics and Preferences, captured demographic and preference-based attitudes. This component described personal background and cultural preferences and exhibited how individuals perceive and interact with insects. The third component (RC3), Emotional and Behavioural Reactions, was characterized by emotional and behavioural responses toward insects. This dimension highlights the affective and instinctual reactions that influence behaviour toward insects.

The Scree Plot (Fig: 1) serves as a visual diagnostic tool that supports the retention of three principal components. It demonstrates a clear “elbow” after the third component, with unrotated eigenvalues of 3.288, 2.603, and 2.077. According to Kaiser’s criterion (eigenvalues > 1), and the cumulative variance explained, three components are justified. The rotated solution further refines interpretability by redistributing variance across components, with slightly adjusted eigenvalues—2.849, 2.658, and 2.461 respectively.

**Figure 1.**
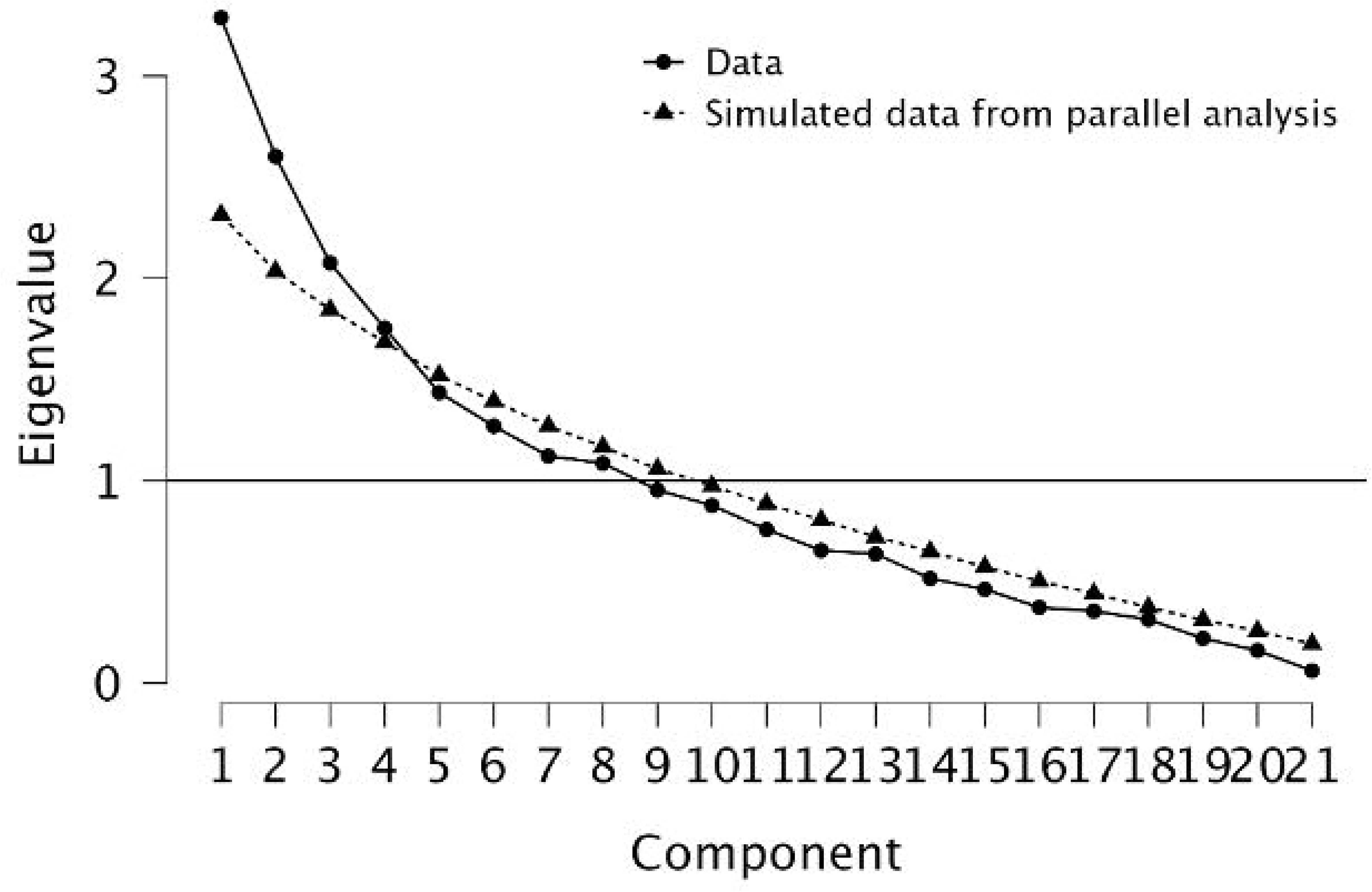
Scree Plot of Principal Components Extracted from Attitudinal Data on Insects.

The Path Diagram (Fig: 2) reflects the structure identified by the PCA, visually mapping the relationships between latent constructs (RC1, RC2, RC3) and observed variables (survey items). Each arrow in the diagram represents the strength and direction of component loadings, with thicker or longer arrows indicating stronger associations. The loadings correspond directly to the PCA output, with RC1 showing particularly high values for environmental knowledge and perception items, such as “Do you consider insects to be helpful to the environment?” (loading = 0.908). RC2 is associated with demographic and preference-related variables (e.g., age, education, and attitudes toward living with insects), while RC3 captures emotional responses—fear, discomfort, and aversion to insects. Notably, the absence of strong arrows linking high-uniqueness items like "Gender" and "Resident" to any component reflects their low communalities and suggests they may represent independent factors not captured within the three-component structure.

**Figure 2.**
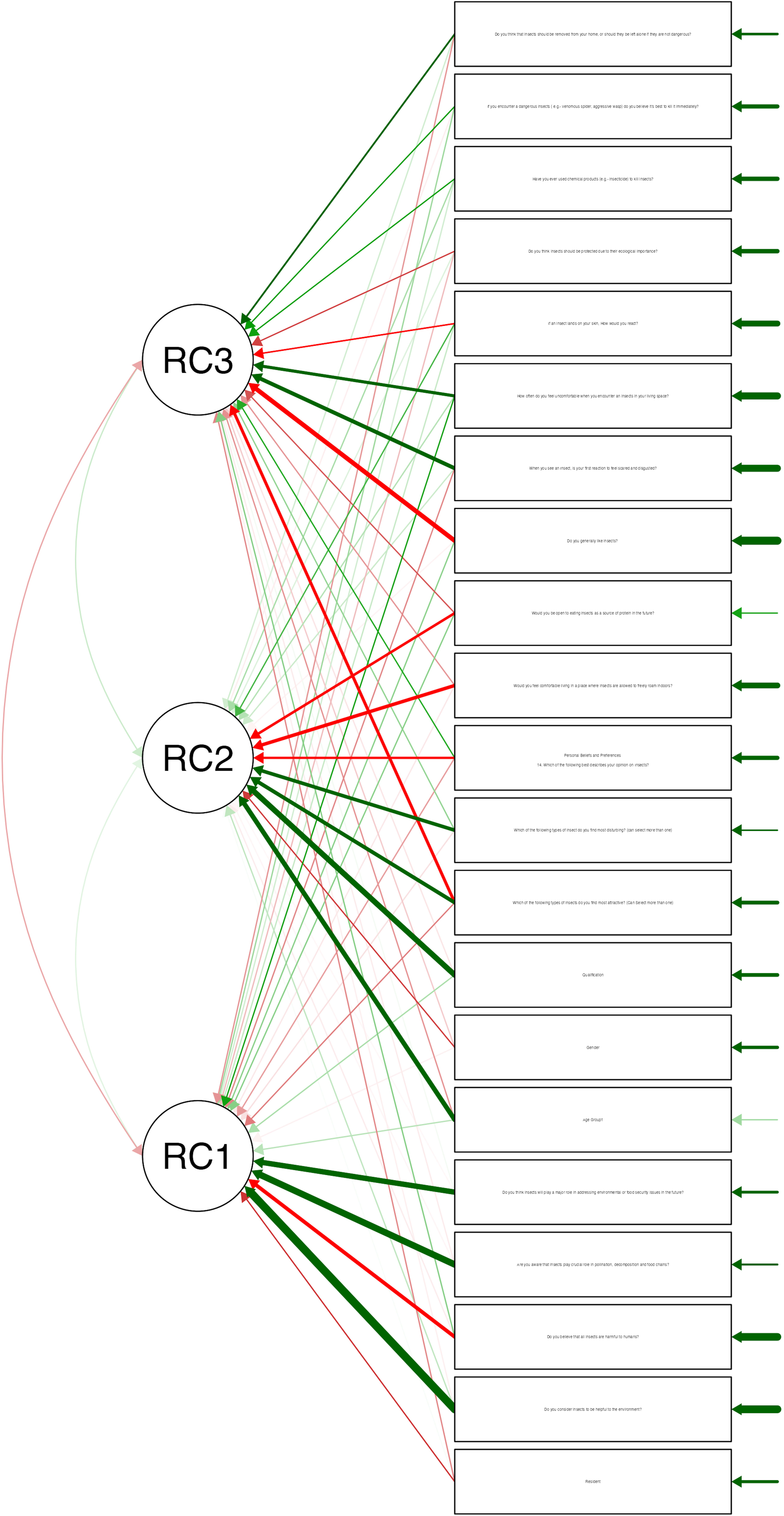
Principal Component Loadings and Interrelations Among Attitudinal Constructs Regarding Insects.

A chi-squared test of model fit yielded a statistically significant result (χ² = 197.476, df = 150, p = 0.006), confirming the presence of structured relationships among the observed variables. However, several variables such as gender, place of residence, beliefs about insect protection, and the use of chemical insecticides exhibited high uniqueness values (above 0.80), indicating that they were not well captured by the extracted components and may reflect independent or culturally specific attitudes. Overall, the PCA revealed a three-dimensional framework— ecological knowledge, demographic influences, and emotional responses—that together shape public perceptions of insects.

## Discussion

From a biological perspective, the paper explores the innate fears humans often harbour towards insects, rooted in evolutionary psychology and reinforced by the complexities of insect social behaviour. These fears can manifest as entomophobia, a specific phobia that disrupts daily life for many individuals. By examining the biological underpinnings of these emotional responses, the discussion reveals how instinctual fears are compounded by cultural attitudes and media portrayals that sensationalize insects, contributing to societal anxieties.

The interplay between attraction and fear is evident in societal attitudes of insects as potential food sources. Entomophagy promotes insect-based diets as sustainable food system alternatives and source of protein. This study addresses the two contrair perceptions about insects, that they are essential components of ecosystems and as sources of concern; this may result in improved and effective educational strategies and conservation practices that respect both ecological integrity and cultural significance.

In conclusion, for most attitudes toward insects are consistent across age groups but are less likely for younger ones to perceive insects as harmful. This indicates an area for intervention, where efforts to educate people about the the ecological significance of insects could counteract negative biases and encourage coexistence. When it comes to gender, females and males have equivalent views about perceptions of insects, except for dealing dangerous insects. Females are likely to prefer immediate removal, indicating a need for targeted public health drives on safe handling and cohabitation. Interestingly, rural and urban residents perceive insects with same attitude, with no statistically significant differences in ecological awareness, attitudes toward their presence, aversion, and usefulness. This suggests that resident status per se does not significantly impact the perception of insects. Education is an important factor in shaping attitudes toward insects. Higher educated people exhibited greater awareness of the ecological role of insects, less fearfulness and discomfort and more tolerant of their presence. Targeted initiatives and life experiences can promote more positive understanding of insects and their dynamic role in our ecosystem.

The survey results indicate interesting paradoxes in public perceptions of insects; a complex relationship between recognition of their ecological value and personal comfort levels were highlighted. These paradoxes highlight the importance of a nuanced understanding of public opinions towards insects. While recognizing the environmental advantages of insects, a significant percentage of the respondents also consider all insects to be dangerous and harmful to humans. Such inconsistency indicates knowledge gap in understanding the beneficial roles of insects in ecosystems. Targeted educational and awareness programs may address such a misconception and help develop a better understanding of the ecological importance of insects. Participants showed a contradictory approach to safeguarding insects based on their ecological significance and the desire to eliminate dangerous insects immediately. This dichotomy may stem from prioritizing safety over ecological concerns in certain circumstances. The survey also revealed agreement that non-dangerous insects should be left alone in the home; contrary to this, there was discomfort with the idea of insects freely moving indoors. This inconsistency could be a balance between wanting control over private space and an awareness of the need to peacefully coexist with harmless insects.

The identified paradoxes highlight the importance of education and awareness campaigns in filling knowledge gaps and promote a more nuanced understanding of insects’ roles in ecosystems.

The results of the principal component analysis reveal strong relationships between variables, as indicated by the chi-squared test. The test identified three underlying factors that collectively explain 38% of the variance in the data. The first component connects demographic variables like qualification and age group to environmental consciousness, suggesting that more educated individuals and older groups tend to appreciate the environmental importance of insects. This relationship is due to education, life experience, and socioeconomic status, that enables individuals with higher qualifications and older age groups being able to better understand the environmental significance of insects. The second factor highlights the relationship between emotional and behavioural responses to insects, including comfort with insects indoors and willingness to have them as food, and personal beliefs and preferences. This suggests that individuals with positive beliefs concerning insects are likely to feel comfortable with them and accept entomophagy.

The third factor addresses on attitudes toward insect conservation and response to harmful insects, revealing that individuals who prefers conservation are less inclined to use chemical pesticides and more wary of harmful insects. These dimensions provide significant information regarding the complex interplay between demographics, environmental knowledge, emotional reactions, and conservation attitudes towards insects and provide a multidimensional understanding of human perceptions and actions concerning insects. Individuals prioritizing conservation are likely driven by environmental concerns and a preference for sustainable approaches.

### Historical Basis

The relationship between humans and insects has been shaped by various cultural and ecological dimensions. Insects are often perceived as inferior life form but still they play a complex role in our ecosystems and cultural narratives. Institutions and Cultural norms can bring better compliance with conservation practices than external sanctions (Infield et al., 2018; Verma et al., 2023). It is crucial in comprehending the role of indigenous communities in biodiversity conservation, as seen in the efforts to integrate the cultural values of the Tongwe people in Tanzania. They leverage their long-term relationships with the land to manage forest sustainability (Infield et al., 2018).

In addition, insects have been integral to several socio-economic systems, affecting from agriculture to culture. Insect folklore, like the grasshopper and the ant fable, narrates moral teachings on diligence and preparedness. These demonstrate how insects have infused human storytelling and cultural norms (Capinera, 2008).

The historical interplay between human societies and insects underscores a larger narrative about our relationship with the nature, mirroring the fluidity and complexity of cultural values that shape our perceptions and behaviours with these. As ecological challenges heighten in the Anthropocene, reconsidering these historic contexts becomes necessary for fostering sustainable and respectful relationships with the myriad forms of life that inhabit on Earth (Cover et al., 2019; Verma et al., 2023).

### Biological Basis

Insects are an important component of ecosystem, and their interactions with other organisms, including plants, demonstrate complex biological mechanisms that shape thoughts about these creatures. For instance, the relationship between insects and plants demonstrate the evolutionary adaptations both groups have developed together. Plants have adapted to defend themselves through resistance mechanisms classified into antibiosis and antixenosis resistance, which protect them from herbivorous insects. Antibiosis defences affect the growth and survival of insects, whereas antixenosis defences control feeding behaviours through structural and chemical factors. (Schowalter, 2022). This dynamic emphasizes how insects are perceived not only as food sources but also as potential threats, influencing human attitudes toward them.

The complexity in social structures of insects may be the reason for a more profound understanding of why humans react to insects with fascination coupled with instinctive fear. Their sensory systems are connected to social behaviour, and their odorant receptor differences influence nest mate recognition and colony dynamics. Metabolic processes such as insulin signalling also affect division of labour in colonies (Singer, 2014).

Moreover, evolutionary psychology posits that the phobia of insects may have evolved as an adaptation to survival. The "food-rejection response of disgust" aligns with a heightened awareness of potential threats, and it is possible that people are evolutionarily predisposed to react fearfully in response to insects because of their connotations with disease and contamination (Gutoskey, 2020; Panjwani, 2023). This notion consistent with studies that are showing that humans exhibit an innate sensitivity to fear-relevant stimuli, which may have been advantageous for survival in ancestral environments where encounters with harmful organisms were more frequent (Panjwani, 2023).

The interwoven interconnection of fear and disgust with respect to insects complicates emotional reaction. Fear is typically understood as a direct response to potential danger; disgust introduces a layer of aversion that can amplify reactions to insects. Studies have shown that people who feel extremely disgusted by spiders also show increased fear, implying that these are not only different but are actually interlinked in how they form impressions of these animals (Lockwood, 2015). Consequently, gaining insight into the biological underpinnings of these interactions gains us an understanding of the general human experience of insects as highlighting the double nature of our reactions—both based on evolutionary past and immediate surroundings.

The fear of insects, known as entomophobia, is a specific phobia that can severely impair the functioning of individuals in their everyday lives. Entomophobia is characterized by an intense and irrational fear of insects, resulting in various emotional, cognitive, and behavioural symptoms (Ashworth, 2023; Jose, 2019). The psychological effect of entomophobia can cause avoidance, forcing individuals to avoid locations where insects may be found, thus interfering with their quality of life (Ashworth, 2023).

It can be attributed to both evolutionary and psychological factors. Humans may possess an innate aversion to certain insects, particularly those that are venomous or vectors, as a survival strategy (Ashworth, 2023). Cognitive biases, increased disgust and anxiety sensitivity, and past traumatic experiences also play a part in the formation and worsening of entomophobia (Ashworth, 2023; Fukano & Soga, 2023).

The entomophobia symptoms can be expressed physically, emotionally, and mentally. Sweating, trembling, palpitations, nausea, and panic attack are some of the physical experiences when confronted by insects (Fukano & Soga, 2023). Overpowering anxiety and fear are experienced emotionally, and there is intrusive thinking and an amplified sense of threat mentally. Diagnosing entomophobia requires a comprehensive evaluation by a psychologist/psychiatrist, that includes clinical interviews and psychological assessments (Jose, 2019). Treatment often involve cognitive behavioural therapy (CBT) and exposure therapy, which assist individuals in gaining resilience and reduce anxiety over time (Mo, 2020; Rowden, 2022).

Perceptions oscillating between anxiety and fascination makes entomophobia a complex phenomenon. Recognizing the cultural values associated with insects can help strengthen community ties and enhance resilience against environmental changes (Bealing & Henegouwen, 2019; Infield et al., 2018). However, conflicts between conservation objectives and community requirements are possible, emphasizing the need to integrate local values into conservation efforts (Verma et al., 2023). Education is central to constructing societal attitudes toward insects, with environmental education programs promoting responsibility and appreciation for the organisms (*Dominant Microorganisms and Insects Shaping Earth’s Ecosystems*, 2025; Duffus et al., 2021).

Investigations into the perceptions of insects have shown how there is an intricate interaction between human attitude and cultural representation. Consumer acceptance of insects as food, or entomophagy, has become a significant area of interest through a scoping study identified that 102 studies reviewed consumer attitudes, preferences, and barriers to insect-based products, emphasising both the potential benefits and challenges to consumer acceptance within different cultural contexts (Tan et al., 2015).

Ruby et al. (2015) reported that 72% of American and 74% of Indian participants were willing to eat insects, with men being more willing than women, particularly in the USA. Disgust was the primary emotional response, although benefits such as nutrition and sustainability were noted, along with concerns of disease. Lemelin et al. (2017) found that North American perceptions range from positive to negative, shaped by socio-cultural contexts, often reflecting taxonomic inaccuracies. Information sources play an important role, and educational strategies are needed to dispel misconceptions.

Diverse attitudes towards insects as food are shown by various comparative studies based on different historical and cultural contexts. Adult participants from various nations highlight the significance of cultural perception in determining attitudes toward eating insects, emphasizing that traditional practices and beliefs greatly influence the acceptance or rejection of insects as a food source (Tan et al., 2015).

The psychological aspects of insect perception have also been in focus. Studies related to entomophobia, demonstrate how disgustingness and anxiety can impact a person’s learning history and self-esteem. For example, research indicates that disgust can be a factor in increasing learning interest in some situations, which may have implications in terms of how individuals learn about insects (Duffus et al., 2021).

The representation of insects in media particularly film and television play a great role in influencing public attitudes. Frequently represented in sensationalized or scary terms, these depictions feed into societal fears and misconceptions about insects. This is important in understanding how media can perpetuate/reinforce entomophobia and affect attitudes toward entomophagy and environmental conservation (“Depictions of Insects in Film and Television,” 2023).

Evolutionary psychology, cultural perception, and media representation all influence the complex human-insect relationship, usually leading to entomophobia. Although humans see the role of insects in the ecosystem as invaluable, there is a widespread inconsistency in opinion, with most regarding them as detrimental. Attitudes are dictated largely by education, with educated individuals tending to be better informed and more tolerant. Focused initiatives can support a more positive perception of insects, filling knowledge gaps and dispelling misconceptions. The tension between fascination and fear characterizes societal orientations toward entomophagy, conservation, and environmental sustainability, exemplifying the importance of empathetic understanding and education in advancing coexistence and appreciation for the critical roles of insects in ecosystems.

## References

Abajue, M. C., & Gbarakoro, T. N. (2023). Prospects of Insect Farming for Food Security, Environmental Sustainability, and as an Alternative to Agrochemical Use. In One Health Implications of Agrochemicals and their Sustainable Alternatives (pp. 565–600). Springer.

Ashworth, R. (2023). The Fear of Insects: Understanding and Overcoming Entomophobia. Medicalnewstoday. https://www.medicalnewstoday.com/articles/entomophobia#summary

Barragán-Fonseca, K. B. (2024). Sustainable Insect Farming: A System and One Health Approach for Effective Management. Animal Science Cases, 2024, ascs20240003.

Bealing, N., & Henegouwen, A. van B. (2019). Bugs: Beauty and Danger. Groundworkgallery. https://www.groundworkgallery.com/exhibition/bugs-beauty-danger/

Berenbaum, M. (1996). Bugs in the system: Insects and their impact on human affairs. Basic Books.

Cadinu, L. A., Barra, P., Torre, F., Delogu, F., & Madau, F. A. (2020). Insect rearing: Potential, challenges, and circularity. Sustainability, 12(11), 4567.

Capinera, J. L. (2008). Encyclopedia of entomology. Springer Science & Business Media.

Cassel, C. S. (2016). Bugs After the Bomb: Insect Representations in Postatomic American Fiction and Film.

Cover, R. M., Strankman, M. A. M., Scott, M. A., & Bundy, C. S. (2019). ENTOMOMANIA: Insects in Art & Culture.

Depictions of Insects in Film and Television,. (2023). ENTOMOPHOBIA GUIDE. https://phobiasolution.com/entomophobia-guide

Dominant Microorganisms and Insects Shaping Earth’s Ecosystems. (2025). https://biologyinsights.com/dominant-microorganisms-and-insects-shaping-earths-ecosystems/

Duffus, N. E., Christie, C. R., & Morimoto, J. (2021). Insect cultural services: How insects have changed our lives and how can we do better for them. Insects, 12(5), 377.

Free, J. B. (1970). Insect pollination of crops.

Fukano, Y., & Soga, M. (2023). Evolutionary psychology of entomophobia and its implications for insect conservation. Current Opinion in Insect Science, 101100.

Gaston, K. J. (1991). The magnitude of global insect species richness. Conservation Biology, 5(3), 283–296.

Gutoskey, E. (2020). Why Are Humans Afraid of Bugs? Mentalfloss. https://www.mentalfloss.com/article/630000/why-do-humans-fear-bugs

Infield, M., Entwistle, A., Anthem, H., Mugisha, A., & Phillips, K. (2018). Reflections on cultural values approaches to conservation: Lessons from 20 years of implementation. Oryx, 52(2), 220–230. Cambridge Core. 10.1017/S0030605317000928

Jose, V. (2019). Why are humans so afraid of insects. Int J Trend Sci Res Dev, 3(2), 2456– 6470.

Lemelin, R. H., Dampier, J., Harper, R., Bowles, R., & Balika, D. (2017). Perceptions of insects: A visual analysis. Society & Animals, 25(6), 553–572.

Lockwood, J. (2015). Fear + Disgust = Entomological Horror. How Insects Complicate Our Aversive Emotions. Psychologytoday. https://www.psychologytoday.com/gb/blog/the-infested-mind/201512/fear-disgust-entomological-horror

Mo. (2020). Feels Like Bugs Are Biting Me? | Unraveling the Mystery. https://wellwisp.com/feels-like-bugs-are-biting-me/

Panjwani, S. (2023). Why are we afraid of bugs? Three theories that explain why crawlies creep us out. Thenationalnews. https://www.thenationalnews.com/lifestyle/wellbeing/2023/12/06/why-are-we-afraid-of-bugs/

Pellmyr, O. (1992). Evolution of insect pollination and angiosperm diversification. Trends in Ecology & Evolution, 7(2), 46–49.

Ramos-Elorduy, J. (2009). Anthropo-entomophagy: Cultures, evolution and sustainability. Entomological Research, 39(5), 271–288.

Rowden, A. (2022). What to know about entomophobia. https://www.medicalnewstoday.com/articles/entomophobia#summary

Ruby, M. B., Rozin, P., & Chan, C. (2015). Determinants of willingness to eat insects in the USA and India. Journal of Insects as Food and Feed, 1(3), 215–226.

Schowalter, T. D. (2022). Insect ecology: An ecosystem approach. Academic press.

Singer, E. (2014). In Bees, a Hunt for Roots of Social Behavior| Quanta Magazine. Accessed September, 2.

Snaddon, J. L., & Turner, E. C. (2007). A child’s eye view of the insect world: Perceptions of insect diversity. Environmental Conservation, 34(1), 33–35.

Soga, M., Gaston, K. J., Fukano, Y., & Evans, M. J. (2023). The vicious cycle of biophobia. Trends in Ecology & Evolution, 38(6), 512–520.

Tan, H. S. G., Fischer, A. R., Tinchan, P., Stieger, M., Steenbekkers, L., & Van Trijp, H. C. (2015). Insects as food: Exploring cultural exposure and individual experience as determinants of acceptance. Food Quality and Preference, 42, 78–89.

Verma, R. C., Waseem, M. A., Sharma, N., Bharathi, K., Singh, S., Anto Rashwin, A., Pandey, S. K., & Singh, B. V. (2023). The role of insects in ecosystems, an in-depth review of entomological research. International Journal of Environment and Climate Change, 13(10), 4340–4348.

Yadav, D., Ray, S., Maheshwari, S., & Saikanth, D. (2023). INFLUENCE OF INSECTS ON ART AND CULTURE. In Recent Trends in Entomology. Academic publication & Distributors, New Delhi.

