## Supplementary Data for "Human Attitudes Toward Insects and Spiders: Exploring the Paradoxes of Ecological Value and Discomfort"

#### Supplementary Data 1 (PDF file link of Google Form used for survey)

https://drive.google.com/file/d/1MZJ9IC-g6TRE6-sHnl837gX9DJxK2FB/view?usp=sharing

**Supplementary Data 2-4, contingency tables**

### 1. Contingency Table by Gender

| Question | Response | Gender | | Total |
|---|---|---|---|---|
| | | Female | Male | |
| 1. Do you generally like insects? | No | 15 | 11 | 26 |
| | Yes | 15 | 16 | 31 |
| | **Total** | 30 | 27 | 57 |
| 2. Do you consider insects to be helpful to the environment? | No | 1 | 2 | 3 |
| | Yes | 29 | 24 | 53 |
| | **Total** | 30 | 26 | 56 |
| 3. When you see an insect, is your first reaction to feel scared and disgusted? | No | 16 | 13 | 29 |
| | Yes | 14 | 14 | 28 |
| | **Total** | 30 | 27 | 57 |
| 4. Do you believe that all insects are harmful to humans? | No | 28 | 25 | 53 |
| | Yes | 2 | 2 | 4 |
| | **Total** | 30 | 27 | 57 |
| 5. How often do you feel uncomfortable when you encounter insects in your living space? | Rarely | 11 | 15 | 26 |
| | Sometimes | 11 | 8 | 19 |
| | Often | 6 | 0 | 6 |
| | Always | 2 | 4 | 6 |
| | **Total** | 30 | 27 | 57 |
| 6. Which of the following types of insects do you find most attractive? (Top 5) | Butterflies | 9 | 10 | 19 |
| | Bees/Wasp, Moths, Butterflies | 3 | 3 | 6 |
| | Bees/Wasp, Ants, Butterflies | 1 | 2 | 3 |
| | Moths, Butterflies | 3 | 1 | 4 |
| | Spiders, Bees/Wasp, Ants, Moths, Butterflies | 2 | 1 | 3 |
| | **Total** | 30 | 26 | 56 |
| 7. Which of the following types of insects do you find most disturbing? (Top 5) | Mosquitoes | 3 | 8 | 11 |
| | Cockroach | 3 | 3 | 6 |
| | Mosquitoes, Cockroach, Spider | 3 | 2 | 5 |
| | Mosquitoes, Cockroach, Ants | 1 | 3 | 4 |

| | | | | |
|---|---|---|---|---|
| | Butterflies | 2 | 1 | 3 |
| | **Total** | 30 | 25 | 55 |
| 8. If an insect lands on your skin, how would you react? | Panic and Scream | 1 | 3 | 4 |
| | Swat it away | 12 | 9 | 21 |
| | Try to gently remove it | 17 | 15 | 32 |
| | **Total** | 30 | 27 | 57 |
| 9. Are you aware that insects play a crucial role in pollination, decomposition, and food chains? | No | 1 | 1 | 2 |
| | Yes | 29 | 25 | 54 |
| | **Total** | 30 | 26 | 56 |
| 10. Do you think insects should be protected due to their ecological importance? | No | 0 | 1 | 1 |
| | Yes | 30 | 25 | 55 |
| | **Total** | 30 | 26 | 56 |
| 11. Have you ever used chemical products (e.g., insecticide) to kill insects? | No | 15 | 14 | 29 |
| | Yes | 15 | 11 | 26 |
| | **Total** | 30 | 25 | 55 |
| 12. If you encounter a dangerous insect, do you believe it's best to kill it immediately? | No | 9 | 17 | 26 |
| | Yes | 21 | 10 | 31 |
| | **Total** | 30 | 27 | 57 |
| 13. Do you think insects should be removed from your home, or left alone if not dangerous? | Left alone | 19 | 13 | 32 |
| | Removed | 11 | 13 | 24 |
| | **Total** | 30 | 26 | 56 |
| 14. Which of the following best describes your opinion on insects? | Beneficial and respected | 14 | 12 | 26 |
| | Mostly harmless, ignored | 11 | 8 | 19 |
| | Pests, should be eliminated | 1 | 0 | 1 |
| | I don't know | 4 | 7 | 11 |
| | **Total** | 30 | 27 | 57 |
| 15. Would you feel comfortable living in a place where insects are allowed to freely roam indoors? | No | 25 | 21 | 46 |
| | Yes | 5 | 6 | 11 |
| | **Total** | 30 | 27 | 57 |
| 16. Would you be open to eating insects as a source of protein in the future? | No | 28 | 21 | 49 |
| | Yes | 2 | 6 | 8 |
| | **Total** | 30 | 27 | 57 |
| 17. Do you think insects will play a major role in addressing environmental or food security issues? | No | 3 | 1 | 4 |
| | Yes | 27 | 26 | 53 |
| | **Total** | 30 | 27 | 57 |

### 2. Contingency Table by Resident Status

| Question | Response | Residency | | Total |
|---|---|---|---|---|
| | | Rural | Urban | |
| 1. Do you generally like insects? | No | 6 | 20 | 26 |
| | Yes | 8 | 23 | 31 |
| | **Total** | 14 | 43 | 57 |
| 2. Do you consider insects to be helpful to the environment? | No | 0 | 3 | 3 |
| | Yes | 14 | 39 | 53 |
| | **Total** | 14 | 42 | 56 |
| 3. When you see an insect, is your first reaction to feel scared and disgusted? | No | 7 | 22 | 29 |
| | Yes | 7 | 21 | 28 |
| | **Total** | 14 | 43 | 57 |
| 4. Do you believe that all insects are harmful to humans? | No | 13 | 40 | 53 |
| | Yes | 1 | 3 | 4 |
| | **Total** | 14 | 43 | 57 |
| 5. How often do you feel uncomfortable when you encounter insects in your living space? | Rarely | 4 | 22 | 26 |
| | Sometimes | 4 | 15 | 19 |
| | Often | 3 | 3 | 6 |
| | Always | 3 | 3 | 6 |
| | **Total** | 14 | 43 | 57 |
| 6. Which of the following types of insects do you find most attractive? (Top 5) | Butterflies | 6 | 13 | 19 |
| | Bees/Wasp, Moths, Butterflies | 2 | 4 | 6 |
| | Bees/Wasp, Ants, Butterflies | 0 | 3 | 3 |
| | Moths, Butterflies | 1 | 3 | 4 |
| | Spiders, Bees/Wasp, Ants, Moths, Butterflies | 0 | 3 | 3 |
| | **Total** | 14 | 42 | 56 |
| 7. Which of the following types of insects do you find most disturbing? (Top 5) | Mosquitoes | 2 | 9 | 11 |
| | Cockroach | 3 | 3 | 6 |
| | Mosquitoes, Cockroach, Spider | 1 | 4 | 5 |
| | Mosquitoes, Cockroach, Ants | 1 | 3 | 4 |
| | Butterflies | 1 | 2 | 3 |
| | **Total** | 14 | 41 | 55 |
| 8. If an insect lands on your skin, how would you react? | Panic and Scream | 1 | 3 | 4 |
| | Swat it away | 5 | 16 | 21 |
| | Try to gently remove it | 8 | 24 | 32 |
| | **Total** | 14 | 43 | 57 |
| 9. Are you aware that insects play a crucial role in pollination, decomposition, and food chains? | No | 0 | 2 | 2 |
| | Yes | 14 | 40 | 54 |
| | **Total** | 14 | 42 | 56 |
| 10. Do you think insects should be protected due to their ecological importance? | No | 0 | 1 | 1 |
| | Yes | 14 | 41 | 55 |
| | **Total** | 14 | 42 | 56 |

| | | | | |
|---|---|---|---|---|
| 11. Have you ever used chemical products (e.g., insecticide) to kill insects? | No | 8 | 21 | 29 |
| | Yes | 6 | 20 | 26 |
| | **Total** | 14 | 41 | 55 |
| 12. If you encounter a dangerous insect, do you believe it's best to kill it immediately? | No | 7 | 19 | 26 |
| | Yes | 7 | 24 | 31 |
| | **Total** | 14 | 43 | 57 |
| 13. Do you think insects should be removed from your home, or left alone if not dangerous? | Left alone | 8 | 24 | 32 |
| | Removed | 6 | 18 | 24 |
| | **Total** | 14 | 42 | 56 |
| 14. Which of the following best describes your opinion on insects? | Beneficial and respected | 6 | 20 | 26 |
| | Mostly harmless, ignored | 6 | 13 | 19 |
| | Pests, should be eliminated | 0 | 1 | 1 |
| | I don't know | 2 | 9 | 11 |
| | **Total** | 14 | 43 | 57 |
| 15. Would you feel comfortable living in a place where insects are allowed to freely roam indoors? | No | 12 | 34 | 46 |
| | Yes | 2 | 9 | 11 |
| | **Total** | 14 | 43 | 57 |
| 16. Would you be open to eating insects as a source of protein in the future? | No | 11 | 38 | 49 |
| | Yes | 3 | 5 | 8 |
| | **Total** | 14 | 43 | 57 |
| 17. Do you think insects will play a major role in addressing environmental or food security issues? | No | 0 | 4 | 4 |
| | Yes | 14 | 39 | 53 |
| | **Total** | 14 | 43 | 57 |

### 3. Contingency Table by Qualification

| Question | Response | Qualification | | | | | | Total |
|---|---|---|---|---|---|---|---|---|
| | | X | XII | Graduation | Post Graduation | PhD | Other | |
| 1. Do you generally like insects? | No | 2 | 4 | 4 | 9 | 7 | 0 | 26 |
| | Yes | 5 | 1 | 6 | 12 | 5 | 1 | 30 |
| | **Total** | 7 | 5 | 10 | 21 | 12 | 1 | 56 |
| 2. Do you consider insects to be helpful to the environment? | No | 0 | 2 | 1 | 0 | 0 | 0 | 3 |
| | Yes | 6 | 3 | 9 | 21 | 12 | 1 | 52 |
| | **Total** | 6 | 5 | 10 | 21 | 12 | 1 | 55 |
| 3. When you see an insect, is your first reaction to feel scared and disgusted? | No | 6 | 1 | 4 | 11 | 6 | 1 | 29 |
| | Yes | 1 | 4 | 6 | 10 | 6 | 0 | 27 |
| | **Total** | 7 | 5 | 10 | 21 | 12 | 1 | 56 |
| 4. Do you believe that all insects are harmful to humans? | No | 7 | 4 | 8 | 21 | 11 | 1 | 52 |
| | Yes | 0 | 1 | 2 | 0 | 1 | 0 | 4 |
| | **Total** | 7 | 5 | 10 | 21 | 12 | 1 | 56 |
| 5. How often do you feel uncomfortable when you encounter insects in your living space? | Rarely | 4 | 2 | 6 | 7 | 5 | 1 | 25 |
| | Sometimes | 2 | 3 | 2 | 9 | 3 | 0 | 19 |
| | Often | 0 | 0 | 2 | 3 | 1 | 0 | 6 |
| | Always | 1 | 0 | 0 | 2 | 3 | 0 | 6 |
| | **Total** | 7 | 5 | 10 | 21 | 12 | 1 | 56 |
| 6. Which of the following types of insects do you find most attractive? (Top 5) | Butterflies | 2 | 1 | 4 | 8 | 3 | 0 | 18 |
| | Bees/Wasp, Moths, Butterflies | 0 | 0 | 0 | 2 | 4 | 0 | 6 |
| | Bees/Wasp, Ants, Butterflies | 0 | 0 | 0 | 1 | 2 | 0 | 3 |
| | Moths, Butterflies | 0 | 0 | 1 | 3 | 0 | 0 | 4 |
| | Spiders, Bees/Wasp, Ants, Moths, Butterflies | 0 | 0 | 0 | 2 | 0 | 1 | 3 |
| | **Total** | 6 | 5 | 10 | 21 | 12 | 1 | 55 |
| 7. Which of the following types of insects do you find most disturbing? (Top 5) | Mosquitoes | 4 | 1 | 2 | 2 | 2 | 0 | 11 |
| | Cockroach | 0 | 1 | 1 | 4 | 0 | 0 | 6 |
| | Mosquitoes, Cockroach, Spider | 0 | 0 | 0 | 3 | 2 | 0 | 5 |
| | Mosquitoes, Cockroach, Ants | 0 | 1 | 0 | 2 | 1 | 0 | 4 |
| | Butterflies | 0 | 0 | 0 | 2 | 1 | 0 | 3 |
| | **Total** | 6 | 5 | 10 | 21 | 12 | 0 | 54 |
| 8. If an insect lands on your skin, how would you react? | Panic and Scream | 0 | 1 | 1 | 1 | 0 | 0 | 3 |
| | Swat it away | 2 | 1 | 3 | 11 | 4 | 0 | 21 |
| | Try to gently remove it | 5 | 3 | 6 | 9 | 8 | 1 | 32 |
| | **Total** | 7 | 5 | 10 | 21 | 12 | 1 | 56 |
| 9. Are you aware that insects play a crucial role in pollination, decomposition, and food chains? | No | 0 | 1 | 1 | 0 | 0 | 0 | 2 |
| | Yes | 6 | 4 | 9 | 21 | 12 | 1 | 53 |
| | **Total** | 6 | 5 | 10 | 21 | 12 | 1 | 55 |
| | No | 0 | 1 | 0 | 0 | 0 | 0 | 1 |
| | Yes | 6 | 4 | 10 | 21 | 12 | 1 | 54 |

| | | | | | | | | |
|---|---|---|---|---|---|---|---|---|
| 10. Do you think insects should be protected due to their ecological importance? | **Total** | 6 | 5 | 10 | 21 | 12 | 1 | 55 |
| 11. Have you ever used chemical products (e.g., insecticide) to kill insects? | No | 4 | 2 | 6 | 12 | 4 | 0 | 28 |
| | Yes | 3 | 2 | 4 | 9 | 8 | 0 | 26 |
| | **Total** | 7 | 4 | 10 | 21 | 12 | 0 | 54 |
| 12. If you encounter a dangerous insect, do you believe it's best to kill it immediately? | No | 2 | 3 | 3 | 8 | 8 | 1 | 25 |
| | Yes | 5 | 2 | 7 | 13 | 4 | 0 | 31 |
| | **Total** | 7 | 5 | 10 | 21 | 12 | 1 | 56 |
| 13. Do you think insects should be removed from your home, or left alone if not dangerous? | Left alone | 5 | 2 | 3 | 14 | 7 | 1 | 32 |
| | Removed | 1 | 3 | 7 | 7 | 5 | 0 | 23 |
| | **Total** | 6 | 5 | 10 | 21 | 12 | 1 | 55 |
| 14. Which of the following best describes your opinion on insects? | Beneficial and respected | 2 | 2 | 3 | 11 | 7 | 1 | 26 |
| | Mostly harmless, ignored | 2 | 0 | 4 | 9 | 3 | 0 | 18 |
| | Pests, should be eliminated | 0 | 0 | 1 | 0 | 0 | 0 | 1 |
| | I don't know | 3 | 3 | 2 | 1 | 2 | 0 | 11 |
| | **Total** | 7 | 5 | 10 | 21 | 12 | 1 | 56 |
| 15. Would you feel comfortable living in a place where insects are allowed to freely roam indoors? | No | 3 | 2 | 9 | 18 | 12 | 1 | 45 |
| | Yes | 4 | 3 | 1 | 3 | 0 | 0 | 11 |
| | **Total** | 7 | 5 | 10 | 21 | 12 | 1 | 56 |
| 16. Would you be open to eating insects as a source of protein in the future? | No | 5 | 4 | 8 | 20 | 11 | 0 | 48 |
| | Yes | 2 | 1 | 2 | 1 | 1 | 1 | 8 |
| | **Total** | 7 | 5 | 10 | 21 | 12 | 1 | 56 |
| 17. Do you think insects will play a major role in addressing environmental or food security issues? | No | 0 | 1 | 1 | 1 | 1 | 0 | 4 |
| | Yes | 7 | 4 | 9 | 20 | 11 | 1 | 52 |
| | **Total** | 7 | 5 | 10 | 21 | 12 | 1 | 56 |

### 4. Contingency Table by Age Group

| Question | Response | Age Group | | | | | Total |
|---|---|---|---|---|---|---|---|
| | | Below 20 | 21–30 | 31–40 | 41–50 | Above 50 | |
| 1. Do you generally like insects? | Yes | 7 | 15 | 1 | 2 | 6 | 31 |
| | No | 6 | 15 | 2 | 1 | 2 | 26 |
| | **Total** | **13** | **30** | **3** | **3** | **8** | **57** |
| 2. Do you consider insects to be helpful to the environment? | Yes | 10 | 29 | 3 | 3 | 8 | 53 |
| | No | 2 | 1 | 0 | 0 | 0 | 3 |
| | **Total** | **12** | **30** | **3** | **3** | **8** | **56** |
| 3. When you see an insect, is your first reaction to feel scared and disgusted? | Yes | 6 | 16 | 2 | 1 | 3 | 28 |
| | No | 7 | 14 | 1 | 2 | 5 | 29 |
| | **Total** | **13** | **30** | **3** | **3** | **8** | **57** |
| 4. Do you believe that all insects are harmful to humans? | Yes | 1 | 2 | 1 | 0 | 0 | 4 |
| | No | 12 | 28 | 2 | 3 | 8 | 53 |
| 5. How often do you feel uncomfortable when you encounter insects in your living space? | Rarely | 7 | 12 | 1 | 2 | 4 | 26 |
| | Sometimes | 5 | 12 | 0 | 0 | 2 | 19 |
| | Often | 0 | 5 | 1 | 0 | 0 | 6 |
| | Always | 1 | 1 | 1 | 1 | 2 | 6 |
| | **Total** | **13** | **30** | **3** | **3** | **10** | **57** |
| 6. Which of the following types of insects do you find most attractive? (Top 5) | Butterflies | 2 | 15 | 1 | 0 | 1 | 19 |
| | Bees/Wasp, Moths, Butterflies | 0 | 3 | 0 | 2 | 1 | 6 |
| | Bees/Wasp | 2 | 0 | 0 | 0 | 1 | 3 |
| | Bees/Wasp, Butterflies | 1 | 1 | 0 | 0 | 1 | 3 |
| | Spiders, Butterflies | 1 | 2 | 0 | 0 | 0 | 3 |
| | **Total** | **6** | **21** | **1** | **2** | **4** | **34** |
| 7. Which of the following types of insects do you find most disturbing? (Top 5) | Mosquitoes | 4 | 5 | 1 | 1 | 0 | 11 |
| | Cockroach | 2 | 3 | 0 | 0 | 1 | 6 |
| | Mosquitoes, Cockroach, Spider | 0 | 2 | 1 | 1 | 1 | 5 |
| | Mosquitoes, Cockroach, Ants | 1 | 2 | 0 | 0 | 1 | 4 |
| | Butterflies | 0 | 2 | 0 | 0 | 1 | 3 |
| | **Total** | **7** | **14** | **2** | **2** | **4** | **29** |
| 8. If an insect lands on your skin, how would you react? | Panic and Scream | 1 | 3 | 0 | 0 | 0 | 4 |
| | Swat it away | 2 | 14 | 2 | 1 | 2 | 21 |
| | Try to gently remove | 10 | 13 | 1 | 2 | 6 | 32 |

| | | | | | | | |
|---|---|---|---|---|---|---|---|
| | **Total** | **13** | **30** | **3** | **3** | **8** | **57** |
| 9. Are you aware that insects play a crucial role in pollination, decomposition, and food chains? | Yes | 11 | 29 | 3 | 3 | 8 | 54 |
| | No | 1 | 1 | 0 | 0 | 0 | 2 |
| | **Total** | **12** | **30** | **3** | **3** | **8** | **56** |
| 10. Do you think insects should be protected due to their ecological importance? | Yes | 11 | 30 | 3 | 3 | 8 | 55 |
| | No | 1 | 0 | 0 | 0 | 0 | 1 |
| | **Total** | **12** | **30** | **3** | **3** | **8** | **56** |
| 11. Have you ever used chemical products (e.g., insecticide) to kill insects? | Yes | 4 | 14 | 2 | 1 | 5 | 26 |
| | No | 7 | 16 | 1 | 2 | 3 | 29 |
| | **Total** | **11** | **30** | **3** | **3** | **8** | **55** |
| 12. If you encounter a dangerous insect, do you believe it's best to kill it immediately? | Yes | 6 | 20 | 1 | 2 | 2 | 31 |
| | No | 7 | 10 | 2 | 1 | 6 | 26 |
| | **Total** | **13** | **30** | **3** | **3** | **8** | **57** |
| 13. Do you think insects should be removed from your home, or left alone if not dangerous? | Removed | 4 | 14 | 1 | 1 | 4 | 24 |
| | Left alone | 8 | 16 | 2 | 2 | 4 | 32 |
| | **Total** | **12** | **30** | **3** | **3** | **8** | **56** |
| 14. Which of the following best describes your opinion on insects? | Beneficial and respected | 6 | 11 | 2 | 1 | 6 | 26 |
| | Mostly harmless and ignored | 2 | 13 | 0 | 2 | 2 | 19 |
| | Pests, should be eliminated | 0 | 1 | 0 | 0 | 0 | 1 |
| | I don't know | 5 | 5 | 1 | 0 | 0 | 11 |
| | **Total** | **13** | **30** | **3** | **3** | **8** | **57** |
| 15. Would you feel comfortable living in a place where insects are allowed to freely roam indoors? | Yes | 6 | 4 | 0 | 0 | 1 | 11 |
| | No | 7 | 26 | 3 | 3 | 7 | 46 |
| | **Total** | **13** | **30** | **3** | **3** | **8** | **57** |
| 16. Would you be open to eating insects as a source of protein in the future? | Yes | 3 | 5 | 0 | 0 | 0 | 8 |
| | No | 10 | 25 | 3 | 3 | 8 | 49 |
| | **Total** | **13** | **30** | **3** | **3** | **8** | **57** |
| 17. Do you think insects will play a major role in addressing environmental or food security issues? | Yes | 12 | 27 | 3 | 3 | 8 | 53 |
| | No | 1 | 3 | 0 | 0 | 0 | 4 |
| | **Total** | **13** | **30** | **3** | **3** | **8** | **57** |
